# Searching for Correlated Conformational Dynamics: Analysis of the NMR Relaxation Dispersions with Akaike's Information Theory and Hierarchical Clustering

**DOI:** 10.1101/176776

**Authors:** Evgenii L. Kovrigin

**Affiliations:** Department of Chemistry, Marquette University, PO Box 1881, Milwaukee, 53201

## Abstract

In this manuscript, I am proposing an approach for identification of correlated exchange in proteins via analysis of the NMR relaxation dispersion data. For a set of spins experiencing exchange, every relaxation dispersion datasets is fit individually and then—globally while paired with every other dataset. The corrected Akaike’ s Information Criteria (AICc) for individual and global fits are used to evaluate the likelihood of two spins to report on the same dynamic event. Application of hierarchical cluster analysis reveals correlated spin groups using the difference in AICcs as a measure of similarity within the pairs. This approach to detection of correlated dynamics is independent of accuracy of best-fit parameters rendering it less sensitive to experimental noise. High throughput and the absence of the operator bias might make it applicable to a relatively lower quality NMR relaxation dispersion data from large and poorly soluble systems.

---

Correlated conformational motions are thought to be involved in a biological function of a number of proteins^*1-8*^. Spin-relaxation Carr-Purcell-Meiboom-Gill (CPMG) NMR experiments provide access to the exchange rates and populations of the nuclear spins undergoing exchange process^*9-12*^. Revealing whether the dynamic events detected at two different sites in the protein structure are *correlated* (that is both sites experience *the same* dynamic event) is a difficult task. Typically, dynamic parameters are determined for individual sites with high precision and then compared to suggest potential correlations (for examples, see studies on RNAse A^*13*^, and Ras GTPase^*8*^). However, such manual analysis is prone to subconscious bias of the researcher: one has to choose which fitting parameter is most important or use some kind of a parameter combination as a ranking factor.

Here I am proposing the approach utilizing Akaike's information theory and statistical clustering for quantitative assessment of similarity between exchange events reported by multiple spins: the Hierarchical Clustering based on Information Theory (HCIT). The HCIT analysis systematically examines all possible pairs of spins and assigns similarity scores to them. Similarity between millisecond-microsecond dynamics of two spins is judged by computing Akaike's Information Criterion^*14-17*^ for the *global* and *individual* fitting of their experimental relaxation dispersions. The *global* model implies that both spins in a given pair are reporting on the same conformational exchange event. Thus, we determine the common exchange rate constant (k_ex_) and populations of states (usually, p_a_ or p_b_ in a two-state model) for both spins while allowing for the individual chemical shift differences and base transverse relaxation rates. The *individual* model implies *independent* conformational exchange events giving rise to relaxation dispersions observed at the two spins. Akaike's Information Theory reveals, based on the values of the corrected Akaike's Information Criterion (AICc), how likely global fitting for the pair of datasets is more correct than individual fitting. The likelihood score is expressed as difference between AICc's for the global and individual fitting results. Using a negative normalized AICc difference as a measure of a distance between sites in each pair, the hierarchical cluster analysis reveals the spins likely to be involved in the same dynamic process in the protein structure. The hypothetical correlations proposed by the hierarchical clustering may be verified experimentally through measurements of activation barriers and isotope effects as well as using sitedirected mutagenesis in proteins.

## Theory

The Hierarchical Cluster analysis based on Information Theory, HCIT, is a rigorous procedure for evaluation of *similarity* between of exchange modes of different spins followed by "blind" grouping of the datasets, without assuming any mechanism of exchange. First step of analysis is to perform pair-wise fitting of spins to establish similarity in pairs. *Individual* fits of two datasets are compared to a *global* fit for the pair using Akaike's Information Criterion. In this treatment, each model (*global* and *individual*) is evaluated on basis of its sum of squares of residuals from fitting and the number of parameters in the equation. The *individual* model for the two-state conformational exchange process includes (1) the exchange rate constant, k_ex_, (2) the population of a major conformer, p_a_, (3) the base relaxation rate, R_2,0_, and (4) the chemical shift difference between the states in exchange, Δω. For two datasets fit individually, the total number of fitting parameters is equal to 8. The *global* model implies common k_ex_ and p_a_ for both spins in the pair still allowing for individual base relaxation rates, R_2,0_, and chemical shift differences between the states, Δω (total of 6 fitting parameters). The Corrected Akaike's Information Criterion, AICc, is computed for both models as^*17*^:

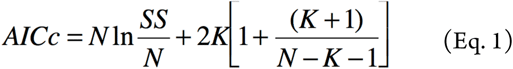

where N is a total number of data points, SS is a sum of squares of residuals, K is the number of fitting parameters plus 1. The model, which has lower AICc, is more likely to be correct. The difference between AICcs may be expressed as evidence ratio, ER, defined as ^*14, 17*^

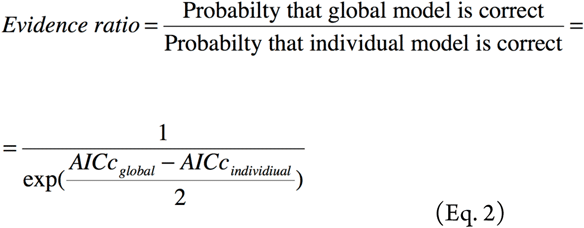

ER value directly reflects how much more likely the global model is correct over the distinct individual models.

The set of N spins produces N*(N-1)/2 possible pairs. To avoid manual comparison of calculated ER values, we subject the results to hierarchical cluster analysis^*18*^. Hierarchical clustering reveals groups, in which spins are better fit by the *global* model when paired to other members of the same cluster, and produce poor global fits when paired to any other spins outside of the cluster. Results of pairwise testing of N datasets are first presented as a resemblance matrix. Figure 1 shows an example resemblance matrix with six datasets. The green and red color is used to indicates relative magnitude of the ER when it is greater than one; crosses stand for ER less than one. According to Eq. 2, values of the ER exceeding one indicate that the global fitting is preferred. In the context of conformational dynamics study, such pair of datasets likely reports on the same dynamic event. In contrast, for spins with ER<1 individual fitting is preferred, implying that these spins reports on distinct conformational dynamics.

**Figure 1.**
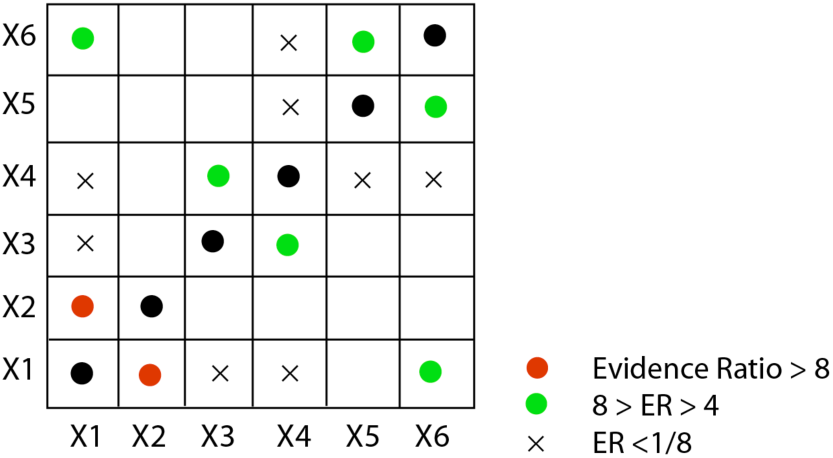
Example resemblance matrix. Each cell of the table is coded according to the value of ER obtained from individual and global fits of each pair of spins. Black circles, diagonal cells not involved in analysis.

Hierarchical clustering is a procedure to help identify nearest neighbors in the set of objects according to a specific measure^*18*^. Using ER as a measure of "distance" between datasets allows to find "closely localized" neighbors that are likely to represent a single dynamic event in the protein structure. Figure 2 shows hierarchical clustering of the dataset from Figure 1, visualizing strong similarity between spins X1 and X2, followed by two other groups X3/X4 and X5/X6. The hierarchical clustering results in a more nuanced grouping than manual analysis because it allows to visualize weaker levels of similarity such as between X1/X2 and X5/X6 clusters based on favorable ER in X1/X6 pair.

**Figure 2.**
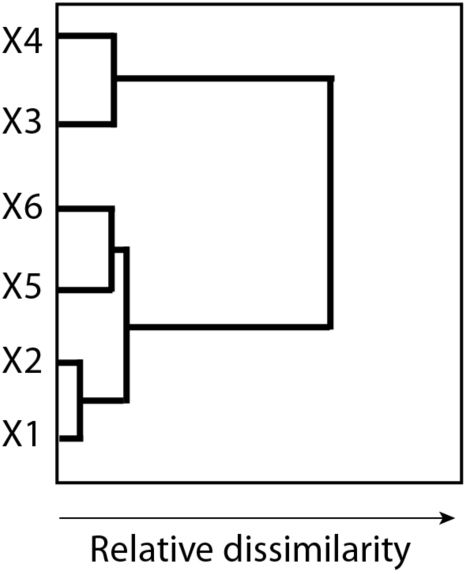
Hierarchical clustering of the six datasets from Figure 1 based on the pairwise ER values.

A crucial distinction of the HCIT analysis from a conventional evaluation based on similarity of fitting results is that the HCIT is performed *without knowledge of best-fit values* of model parameters. Instead, only the sum of squares and total number of parameters are involved in calculations, which renders the entire procedure independent of accuracy of the fit parameters as long as the best-fit curve is successfully produced. The anticipated downside of this treatment is that the low-quality dataset may be successfully clustered with *nearly any other* datasets because its noise level renders SS of the best fit less sensitive to the model type. Therefore, for success of the HCIT, one should avoid inclusion of datasets with a relatively low signal-to-noise ratios.

## Results

To test robustness of the HCIT analysis of the relaxation dispersion data, I simulated relaxation datasets for a number of spins reporting on three distinct exchange events as well as spins that report additional individual dynamics (Table 1). Figure 3 visualizes parameters of these dynamic groups and uncorrelated spins.

**Table 1.** Two-state exchange parameters and chemical shift differences utilized in simulations of relaxation dispersion data.

| Spin # | $k_{ex}$, s<sup>-1</sup> | $p_a$ | $\Delta\omega$, s<sup>-1</sup> |
| --- | --- | --- | --- |
| Fast group |  |  |  |
| 1, 2, 3, 4 | 900 | 0.95 | 100, 200, 300, 400 |
| Slow group |  |  |  |
| 5, 6, 7, 8 | 100 | 0.65 | 100, 200, 300, 400 |
| Intermediate group |  |  |  |
| 9, 10, 11, 12 | 500 | 0.80 | 100, 200, 300, 400 |
| Uncorrelated spins |  |  |  |
| 13 | 100 | 0.95 | 100 |
| 14 | 500 | 0.9 | 400 |
| 15 | 800 | 0.8 | 200 |
| 16 | 1300 | 0.7 | 300 |

**Figure 3.**
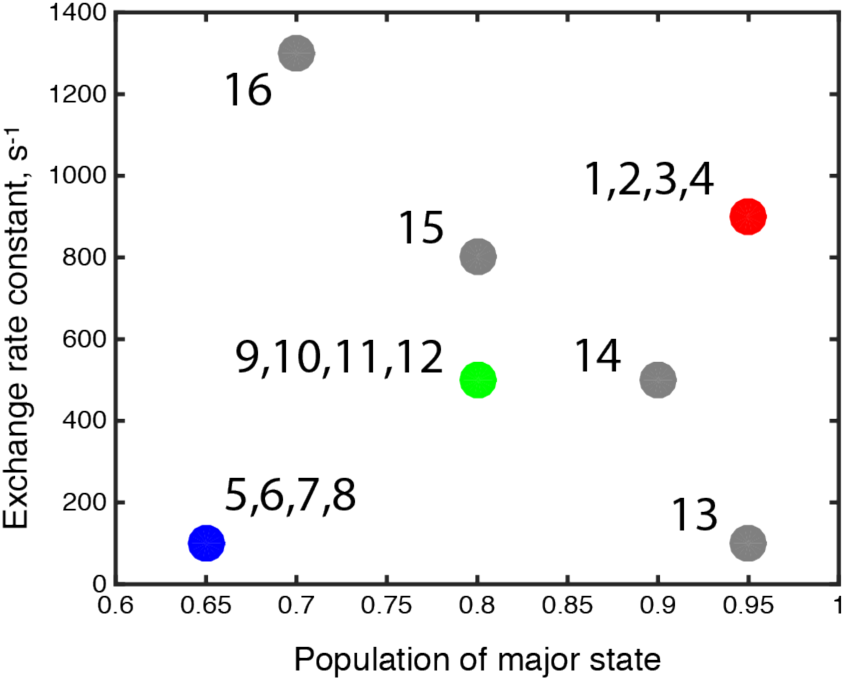
Exchange process parameters used for simulations. Fast group, red; intermediate group, green; slow group, blue; uncorrelated spins; gray. Spin numbers are indicated for each group.

Using these exchange parameters and a typical set of CPMG frequencies (Table 2) with the two-site all-timescale relaxation dispersion equation^*19*^, I simulated the transverse relaxation rate constants, R_2_, for the all spins in Table 1. Supporting Figures 1-4 visualize the corresponding relaxation dispersion profiles. It is important to note, that "shared dynamics" in the spin groups means the same k_ex_ and populations of the two states, which impacts the shape of the curve. The spins may have distinct individual Δω values leading to different amplitude of the dispersion. The obtained R_2_ values were further used for calculation of peak intensities thus simulating outcome of a standard rcCPMG experiment^*20*^ performed in triplicate with a two-point rate determination (for details see O'Connor and Kovrigin^*8*^).

**Table 2.** CPMG frequencies and the total relaxation time.

| # | 1 | 2 | 3 | 4 | 5 | 6 |
| --- | --- | --- | --- | --- | --- | --- |
| $\nu$, Hz | 50 | 75 | 100 | 125 | 150 | 200 |
| T, ms | 40 | 26.7 | 40 | 32 | 40 | 40 |
| # | 7 | 8 | 9 | 10 | 11 |  |
| $\nu$, Hz | 250 | 300 | 350 | 400 | 520 | |
| T, ms | 40 | 40 | 40 | 40 | 38.5 |  |

To assess sensitivity of HCIT analysis to the signal-to-noise ratio in the data, different amounts of random noise were applied to peak intensities to simulate different relative acquisition times (Table 3). The obtained datasets were subject to standard fitting procedures^*8*^ both individually or globally in pairs. The obtained SS along with number of parameters in the fits were subject to hierarchical clustering based on the ER computed for the individual and pairwise fits. All in-house code necessary for simulations and analysis was written in MATLAB and available upon request.

**Table 3.** Addition of noise to the simulated data.

| Dataset quality | High (ideal) | Medium (good practical) | Low (typical) |
| --- | --- | --- | --- |
| Signal/noise, S/N | 1000 | 180 | 128 |
| Acquisition time equivalents | 1 | 1/30 | 1/60 |

Figure 4 demonstrates output of the HCIT analysis in case when signal-to-noise ratio, S/N, is very high (1000). Spin names are listed along the vertical axis. Original group of each spin is indicated with a horizontal bar on the left with the same color code as in Figure 3. As anticipated, all spins from the groups are gathered in corresponding clusters. The "uncorrelated" spins X13 and X16 are not closely grouped with any other due to very distinct dynamic modes. Spins X15 and X14 are joined to the fast and intermediate groups, respectively. This grouping reflects stronger effect of k_ex_ on the clustering results than the population difference, which may be rationalized recalling that k_ex_ value affects the *shape* of the relaxation dispersion curve, while p_a_ is mostly responsible for the amplitude of the dispersion (R_ex_).

**Figure 4.**
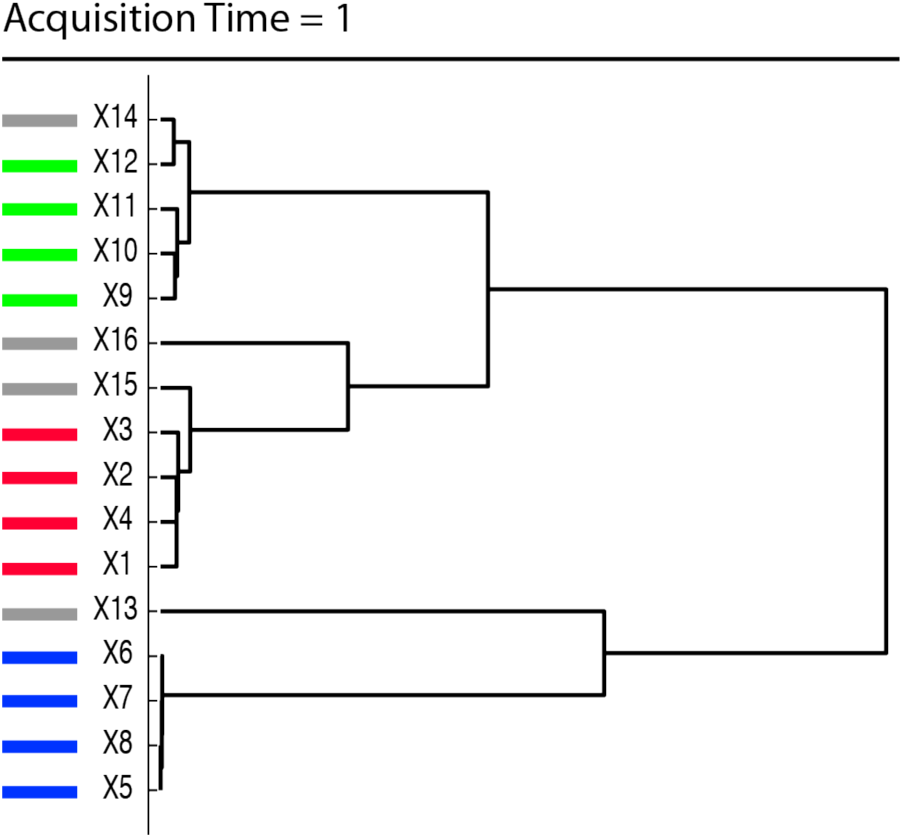
Hierarchical clustering of spins according the ER values for the global vs. individual fits at a very high signal-to-noise ratio (S/N=1000).

To simulate data with the S/N of a feasible relaxation dispersion measurement^*8*^, we need to shorten the acquisition time by a factor of 30 (S/N decreases to 180). Due to increasing noise in the peak intensities, accuracy of the fitted R_2_ decreases. As a result, dispersion curves of the spins with smallest chemical shift differences become poorly defined, making us to remove them from analysis (X1 and X9). Figure 5 demonstrates HCIT analysis of the remaining sensitive datasets. The correlated spins mostly remain grouped together but groups are now more significantly "contaminated" with uncorrelated spins. Slow exchange group (blue) remains most stable and this is correlated with its greatest separation from others in k_ex_ vs. p_a_ graph in Figure 3.

**Figure 5.**
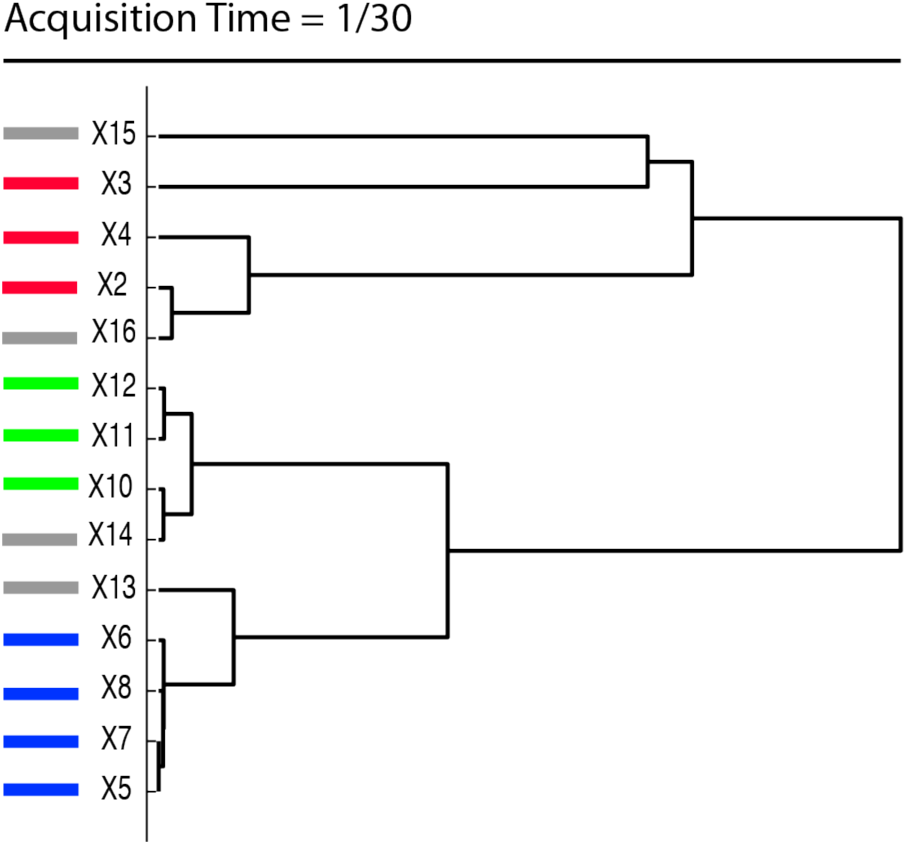
Hierarchical clustering of spins according the ER values for the global vs. individual fits at a practical signal/noise ratio (S/N=180). Acquisition times of this "experiment" is 1/30 of the one in Figure 4. The datasets X1, and X9 with R_ex_ smaller than 4x R_2_ RMSD were removed from analysis.

If the acquisition time is further cut by a factor of two, we have to remove additional datasets, X5 and X13, due to errors in R_2_ becoming comparable to the amplitude of their relaxation dispersion curves. The HCIT analysis still correctly keeps the grouped spins together, though allows the inclusion of the X14 (uncorrelated but close in Figure 3) into the intermediate group (green) before X11 is clustered.

**Figure 6.**
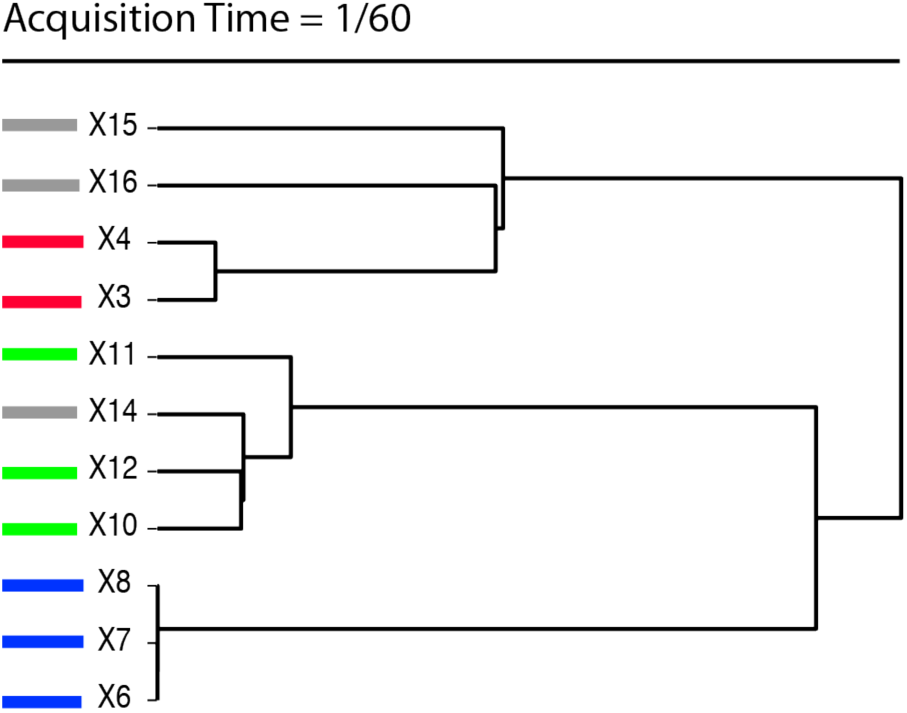
Hierarchical clustering of spins according the ER values for the global vs. individual fits at S/N=128, corresponding to one half of the acquisition time of the "experiment" in Figure 5. The datasets X1, X5, X9, and X13 with R_ex_ smaller than 4x R_2_ RMSD were removed from analysis.

## Discussion

The analysis of simulated data demonstrates how HCIT allows for automated grouping of spins without consideration of the values of particular fitting parameters. In a sense, it compares *shapes* of the relaxation dispersion profiles and will work even if the fitting procedure is not expected to produce very accurate best-fit parameters. This is particularly important because the Carver-Richards equation^*19*^ in the intermediate and fast exchange regimes was shown to provide unstable fits to the relaxation dispersion data obtained at a single static magnetic field^*21*^. The data discussed in this manuscript were simulated using one static magnetic field value. Future simulations with two static magnetic fields will help determine whether HCIT algorithm provides a meaningful degree of compensation to the loss of sensitivity in a one-field setting. In other words, even if the fitting parameters may not be determined at one magnetic field, HCIT might be able to reliably detect correlated spin groups in such datasets. In many studies, quantitative parameters of the exchange dynamics in proteins are not as important by themselves as reliable *mapping of distinct dynamic modes* on the protein structure. Ability to perform such qualitative analysis may be a key to extending reach of spin relaxation NMR measurements to large and challenging systems, where the high S/N value is not practically achievable.

## Acknowledgement

The author is indebted to Dr. James Kempf and Dr. Anthony Mittermaier for helpful discussions of this material.

## Supporting Information

**Supporting Figure S1.**
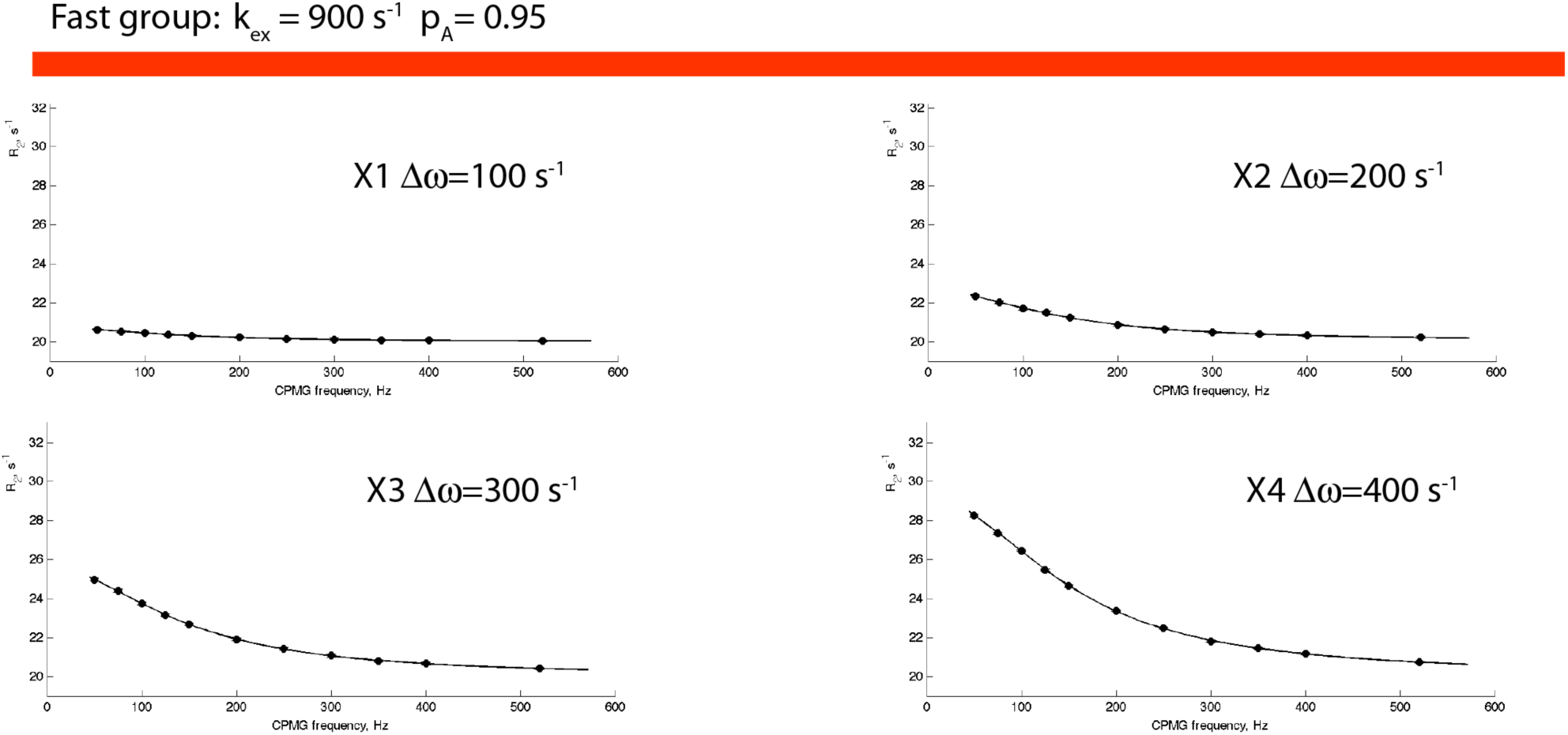
Simulated relaxation dispersion curves for the "fast" spin group in Table 1. The horizontal bar is color coded according to Figure 3 for easier reference.

**Supporting Figure S2.**
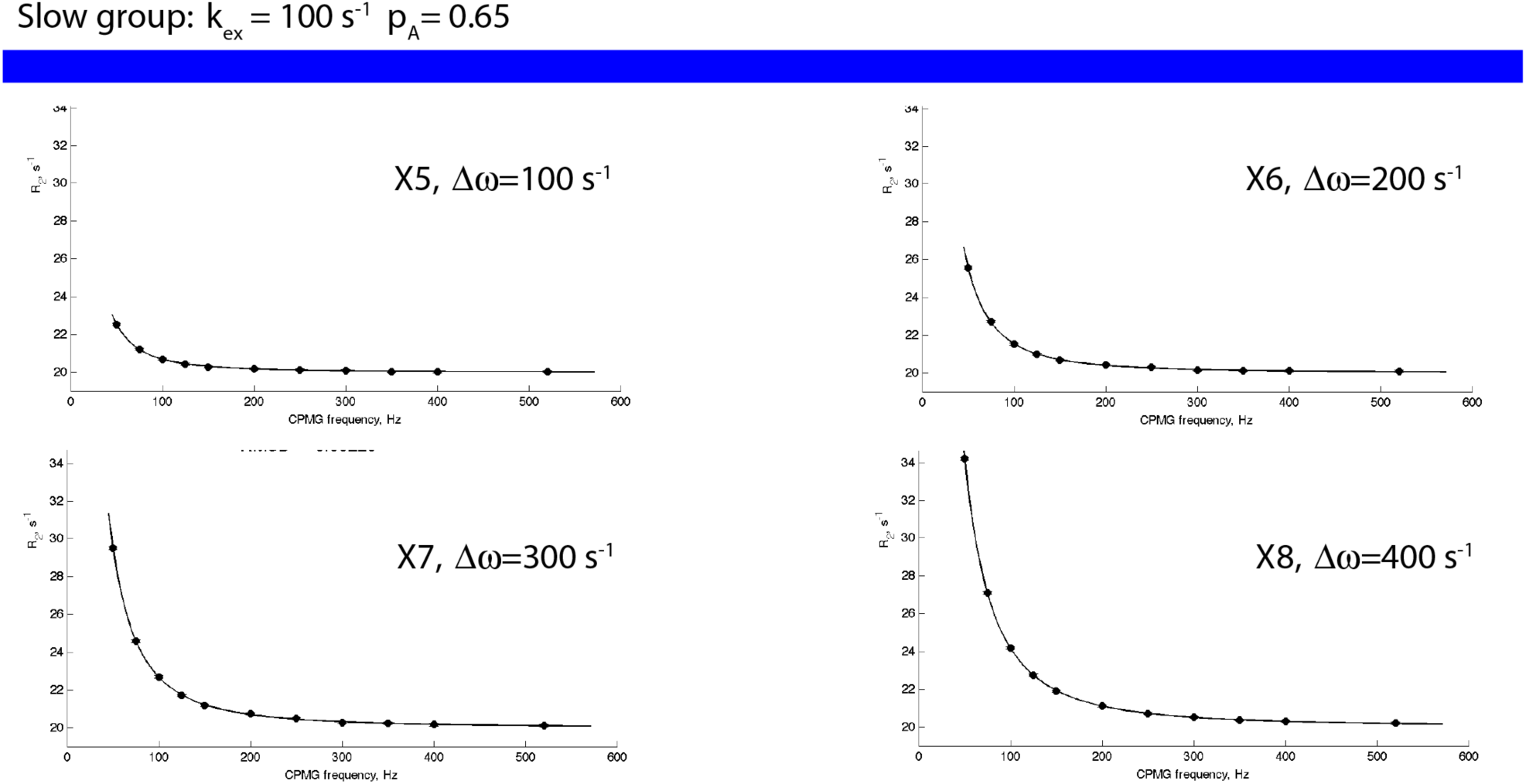
Simulated relaxation dispersion curves for the "slow" spin group in Table 1.. The horizontal bar is color coded according to Figure 3 for easier reference.

**Supporting Figure S3.**
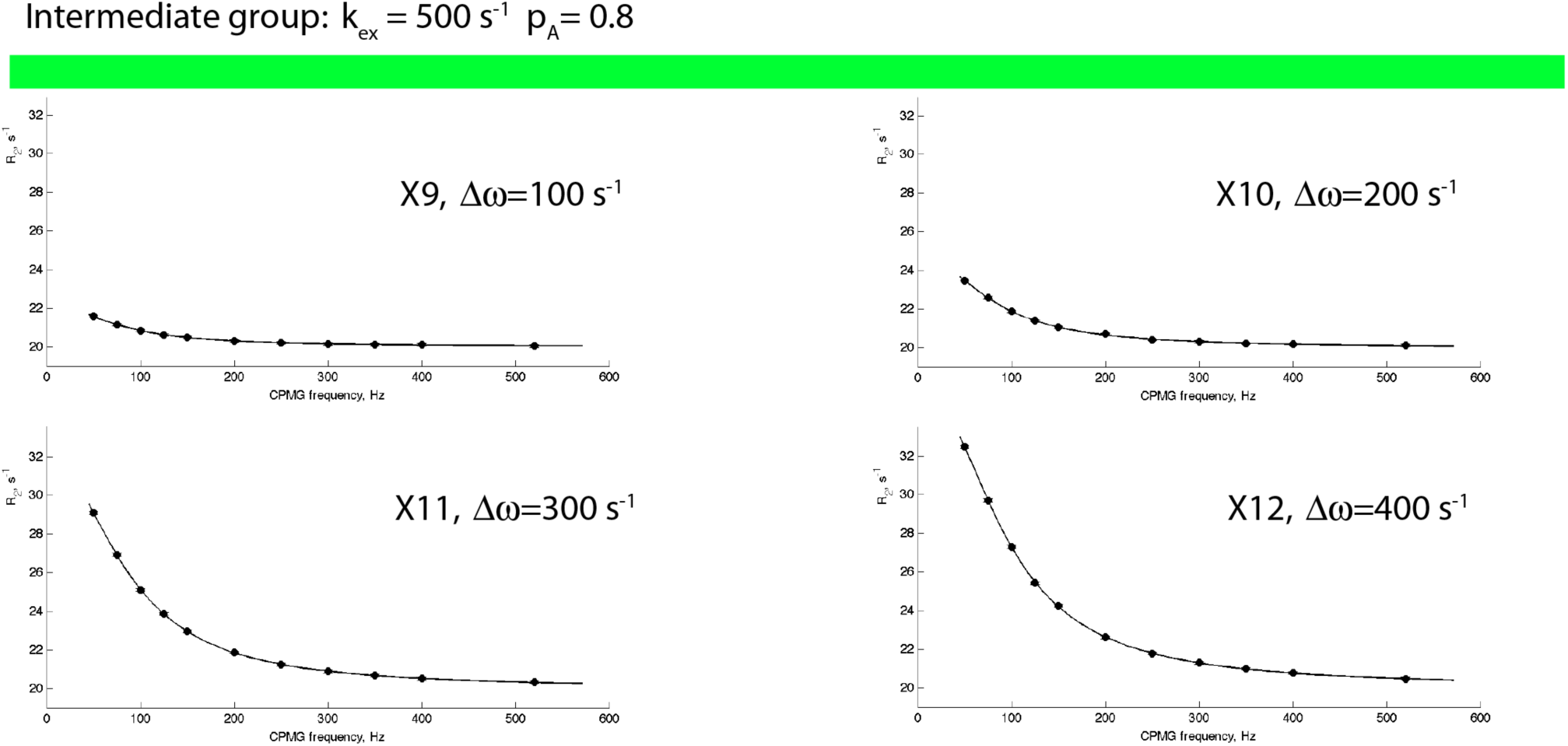
Simulated relaxation dispersion curves for the "intermediate" spin group in Table 1. The horizontal bar is color coded according to Figure 3 for easier reference.

**Supporting Figure S4.**
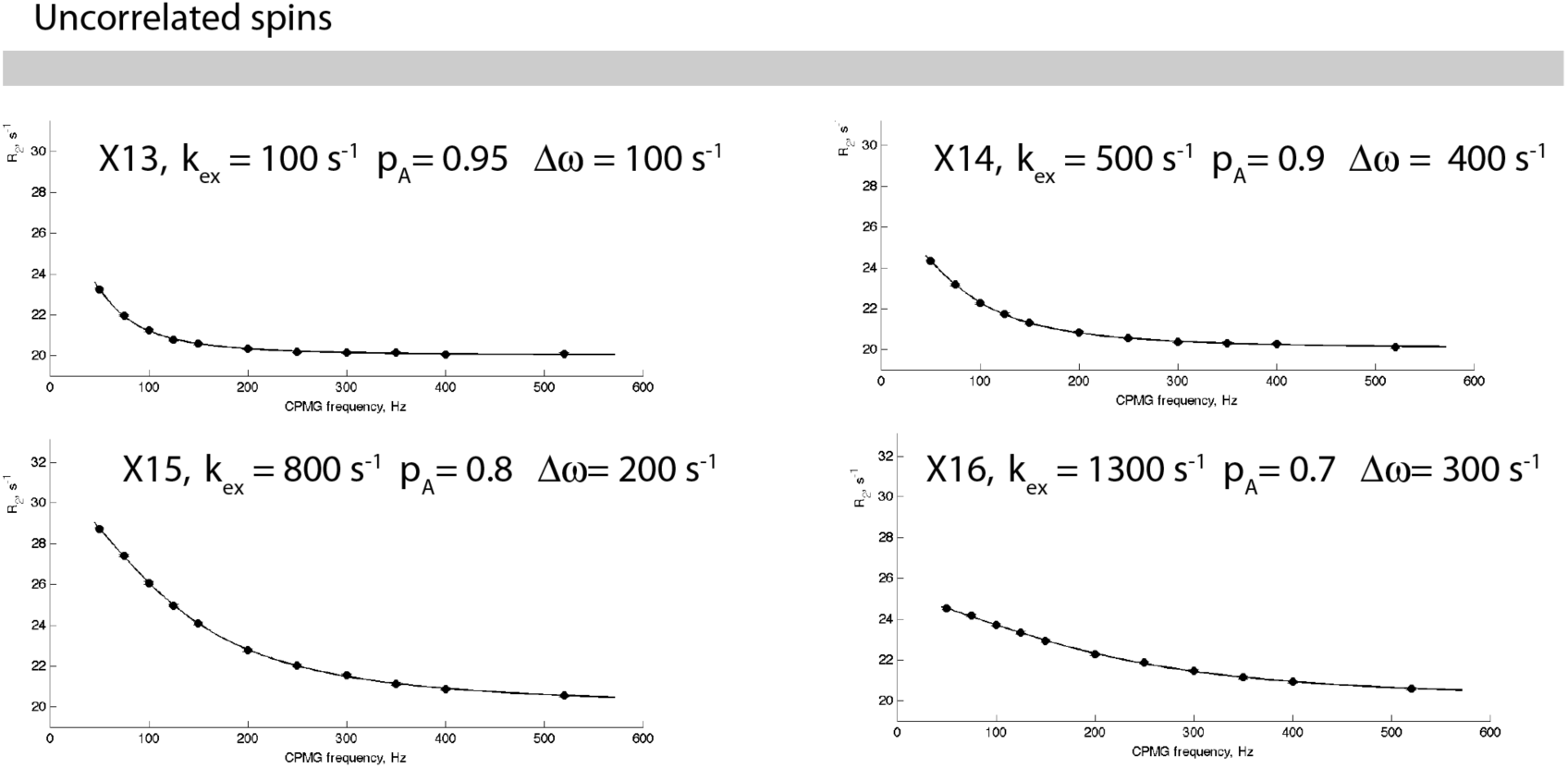
Simulated relaxation dispersion curves for the uncorrelated spins in Table 1. The horizontal bar is color coded according to Figure 3 for easier reference.

## References

[1] Mueller, G. A., Pari, K., DeRose, E. F., Kirby, T. W., and London, R. E. (2004) Backbone dynamics of the RNase H domain of HIV-1 reverse transcriptase, Biochemistry 43, 9332–9342.

[2] Henzler-Wildman, K., and Kern, D. (2007) Dynamic personalities of proteins, Nature 450, 964–972.

[3] Eisenmesser, E. Z., Millet, O., Labeikovsky, W., Korzhnev, D. M., Wolf-Watz, M., Bosco, D. A., Skalicky, J. J., Kay, L. E., and Kern, D. (2005) Intrinsic dynamics of an enzyme underlies catalysis, Nature 438, 117–121.

[4] Volkman, B. F., Lipson, D., Wemmer, D. E., and Kern, D. (2001) Two-state allosteric behavior in a single-domain signaling protein, Science 291, 2429–2433.

[5] Manley, G., and Loria, J. P. (2012) NMR insights into protein allostery, Archives of Biochemistry and Biophysics 519, 223–231.

[6] Lipchock, J. M., and Loria, J. P. (2010) Nanometer Propagation of Millisecond Motions in V-Type Allostery, Structure 18, 1596–1607.

[7] Kovrigin, E. L., and Loria, J. P. (2006) Enzyme dynamics along the reaction coordinate: Critical role of a conserved residue, Biochemistry 45, 2636–2647.

[8] O'Connor, C., and Kovrigin, E. L. (2008) Global conformational dynamics in Ras, Biochemistry 47, 10244–10246.

[9] Carr, H. Y., and Purcell, E. M. (1954) Phys. Rev. 94, 630–638.

[10] Meiboom, S., and Gill, D. (1958) Rev. Sci. Instrum. 29, 688–691.

[11] Palmer, A. G., Kroenke, C. D., and Loria, J. P. (2001) Nuclear magnetic resonance methods for quantifying microsecond-to-millisecond motions in biological macromolecules, Methods in Enzymology 339, 204–238.

[12] Kay, L. E. (1998) Protein dynamics from NMR, Biochemistry and Cell Biology-Biochimie Et Biologie Cellulaire 76, 145–152.

[13] Cole, R., and Loria, J. P. (2002) Evidence for flexibility in the function of ribonuclease A, Biochemistry 41, 6072–6081.

[14] Akaike, H. (1973) Information theory and an extension of the maximum likelihood principle, In *Second International Symposium on Information Theory* (Petrov, B. N., and Csaki, B. F., Eds.), pp 267–281, Academiai Kiado, Budapest.

[15] Akaike, H. (1981) Likelihood of a model and information criteria, Journal of Econometrics 16, 3–14.

[16] Bozdogan, H. (1987) Model selection and Akaike's Information Criterion (AIC): The general theory and its analytical extensions, Psychometrika 52, 345–370.

[17] Motulsky, H., and Christopoulos, A. (2004) Fitting Models to Biological Data Using Linear and Nonlinear Regression: A Practical Guide to Curve Fitting, 1st ed., Oxford University Press, USA.

[18] Romesburg, C. (2004) Cluster Analysis for Researchers, Lulu.com.

[19] Carver, J. P., and Richards, R. E. (1972) A general two-site solution for the chemical exchange produced dependence of T2 upon the Carr-Purcell pulse separation, J. Magn. Reson. 6, 89–105.

[20] Loria, J. P., Rance, M., and Palmer, A. G. (1999) A relaxation-compensated Carr-Purcell-Meiboom-Gill sequence for characterizing chemical exchange by NMR spectroscopy, Journal of the American Chemical Society 121, 2331–2332.

[21] Kovrigin, E. L., Kempf, J. G., Grey, M. J., and Loria, J. P. (2006) Faithful estimation of dynamics parameters from CPMG relaxation dispersion measurements, Journal of Magnetic Resonance 180, 93–104.

